# Safety learning produces rapid fear suppression and distinct amygdala–prefrontal engagement

**DOI:** 10.64898/2026.08.06.743388

**Authors:** Maimoona Altaf, Chulmin Cho, Teresa Ann Maletta, Sung Lim, Loren J. Martin, Hugo Lehmann, Neil M. Fournier

## Abstract

Animals detect and evaluate signs of danger and safety in their environment to ensure survival, yet the neural mechanisms that distinguish safety learning from other forms of conditioned inhibition, remain poorly understood. Here, we directly compared fear and safety learning in male rats. Fear conditioned rats showed high freezing to the tone and the conditioning context, whereas safety conditioned rats showed significant tone-specific reduction in freezing. This safety cue could also generalize to a novel, previously unassociated threat context leading to suppressed freezing when presented demonstrating that inhibitory actions of safety cues are not tied to its original training environment but can modify fear expression across settings. Fear and safety learning also produced unique patterns of neuronal activation and glutamatergic receptor expression in the medial prefrontal cortex (mPFC), basolateral amygdala (BLA), and central amygdala (CeA), as measured by c-Fos immunohistochemistry and Western blotting. Fear conditioning induced greater Fos expression in the BLA and CeA, as well as elevated amygdalar NMDA receptor (GluN1) levels, whereas safety learning increased amygdalar PSD-95 and AMPA receptor (GluA1) expression. Both safety and fear learning increased mPFC Fos expression without affecting glutamatergic receptors levels. Finally, safety conditioning was associated with lower tone-evoked freezing than fear conditioned rats across early extinction sessions and was accompanied by distinct patterns of prefrontal and amygdala activation across extinction. Together, these findings suggest that safety learning engages neural and behavioral mechanisms distinct from fear learning and extinction, while modifying amygdala-prefrontal circuits towards more rapid fear suppression.

## 1. Introduction

The ability to identify dangerous and imminent threats and rapidly initiate defensive responses is critical for survival across species [1,2]. Although the adaptative value of such behaviors is clear, it is also equally imperative that defensive behaviors are suppressed once sources of threat have passed [2]. Understanding how fear responses can be inhibited in situations where they are no longer necessary or have become maladaptive has been the focus of extensive research.

To date, most of this work has examined neural and behavioral mechanisms underlying fear extinction. During fear extinction, defensive responses in rodents (i.e. increased freezing, suppressed grooming, diminished exploration, and avoidance) to a stimulus that was previously predictive of threat becomes gradually diminished through the repeated presentation of the same stimulus in the absence of an aversive outcome. However, another form of conditioned inhibition is known as safety learning [3]. Unlike extinction, safety learning refers to processes by which defensive responses become diminished by associating cues that explicitly signal the absence of threat [4–10]. In this form of inhibitory learning, the presentation of the safety cues serves as a powerful predicator of the non-occurrence or omission of the aversive event by signaling that the animal is “safe” and free from danger, even in situations that might otherwise anticipate threat. Indeed, safety learning has been shown to be highly effective in reducing defensive responses to fearful stimuli in across a range of animals, including rodents, nonhuman primates, and adult humans [3,7].

Many studies have highlighted the amygdala as a key node in threat learning [11,12]. Within the amygdala, the basolateral amygdala (BLA) serves as a major convergence site for integrating sensory inputs with aversive stimuli important for acquiring and recalling cue-threat associations [13–15], while the central amygdala (CeA) and its outputs are critical in the expression of behavioral and physiological responses to learned fear [16,17]. However, growing evidence also indicates that the amygdala not only contributes to threat learning but also actively participates in some forms of inhibitory learning, including extinction and safety learning [3,7,9,18]. Beyond the amygdala, the medial prefrontal cortex (mPFC) also plays a critical role in extinction and safety learning by exerting top-down control over amygdala activity necessary for suppressing fear [12,19–22]. Together, these regions form an integrated circuit, in which the amygdala and mPFC encode and evaluate cues associated with danger and safety.

While the above findings suggest that extinction and safety learning can reduce learned threat responses, they may do so through distinct neural circuits and mechanisms. In the present study, we tested this by directly comparing rats that underwent fear (explicitly paired tone-shock) and safety (explicitly unpaired tone-shock) conditioning to determine whether prior safety learning would alter subsequent extinction of conditioned fear. Because the neural circuitry that dissociates safety from extinction learning remains poorly understood, we examined activation and molecular changes in the mPFC and the amygdala to gain further insight into their contribution to fear and safety learning. Importantly, deficits in safety learning have been implicated in anxiety disorders and post-traumatic stress disorder (PTSD), where safety signals fail to reduce threat responses leading to inappropriate generalization of fear [23–27]. By better understanding how safety signals inhibit threat circuits, it may be possible to identify novel therapeutic strategies designed to enhance safety learning and improve fear inhibition in clinical populations.

## 2. Results

### Safety Cues Suppress Conditioned Fear

Rats underwent three days of fear (n=13) or safety (n=17) conditioning. During fear conditioning, a tone stimulus (conditioned stimulus; CS) co-terminated with the footshock (unconditioned stimulus; US). In contrast, during safety conditioning, tone stimuli and footshocks were explicitly unpaired, with the footshocks delivered at pseudorandom intervals between tone presentations. This ensured that the tone reliably signaled for the absence of the shock. As a control, a tone-only group (n=4) was exposed to the same number of tones without any footshocks.

We analyzed the average the percentage of freezing during presentations of the tones across each day of conditioning. By Day 2 of conditioning, the fear conditioned group displayed higher average tone-evoked freezing than the safety and tone-only groups [ps<.012] with these group differences further increasing by Day 3 [p<.001, **Supplementary Fig. S1A**]. Only the fear conditioned group showed a significant increase in average tone freezing across the three conditioning sessions [paired t-tests, ps<.001]. These differences were also evident when analyzing the individual CS (tone) presentations during each conditioning session with tone-evoked freezing increasing gradually for the fear conditioned group than for the safety and tone-only groups [**Supplementary Fig. S1B**]. However, despite differences in tone-evoked freezing, the fear and safety conditioned groups displayed similar levels of post-shock freezing across the three conditioning sessions [**Supplementary Fig. S1C**]. Importantly, both groups showed a significant increase in post-shock freezing from Day 1 to Day 2 of conditioning [**Supplementary Fig. S1C**].

To assess that the rats had successfully acquired appropriate cue-specific fear and safety associations, we performed a recall test twenty-four hours after the last conditioning session. During recall, all rats were exposed to the same context used for conditioning and were presented with three test tone CSs. A 3 x 3 x 2 mixed design ANOVA with Group (fear vs. safety vs. tone-only) as the between subject factor and Tone (1-3) and Interval (preCS vs. CS) as the within-subject factors revealed a significant Group × Interval interaction [F(2,31)=10.01, p<.001, η² =.39], along with significant main effects of Group [F(2,31)=5.42, p=.010, η² =.26] and Tone [F(2,62)=8.51, p<.001, η² =.22]. Follow-up comparisons showed that fear conditioned rats displayed higher freezing during tone CSs than during the preCS periods, particularly during Blocks 1 and 3 [all ps < .001; **Fig. 1A**]. In contrast, safety conditioned rats showed the opposite pattern demonstrating less freezing during tone CS presentations than during the corresponding preCS periods with this effect being most apparent during Blocks 1 and 2 [RM-ANOVA, ps < .010; **Fig. 1B**]. The tone-only rats displayed consistently low and stable levels of freezing across both preCS and CS periods [**Fig. 1C**].

**Fig. 1.**
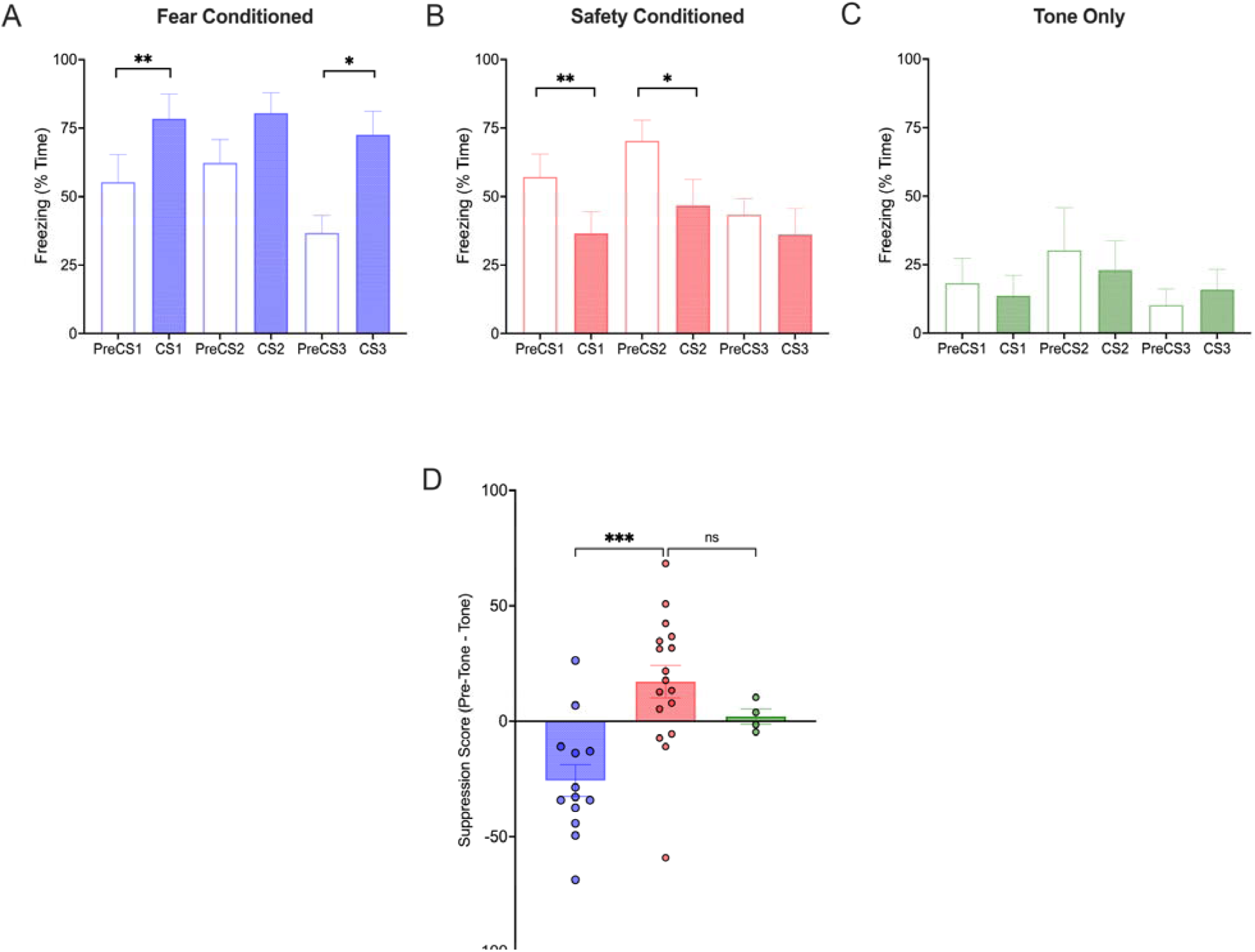
Safety cues suppress freezing during recall. Twenty-four hours after the final conditioning session, rats were returned to the conditioning context and presented with three tone conditioned stimuli (CSs). Freezing was measured during each 20 s CS and the immediately preceding 20 s tone-free (PreCS) period. (A) Fear conditioned rats showed greater freezing during CS presentations than during the PreCS periods. (B) Safety conditioned rats showed reduced freezing during CS presentations relative to the PreCS periods, consistent with the tone functioning as a learned safety signal. (C) Tone-only control rats showed low and stable freezing across the PreCS and CS periods. (D) Group means and individual suppression scores. Positive values indicate lower freezing during the safety cue (CS) than during the tone-free (preCS) period, whereas negative values indicate greater freezing during the tone (CS) than during the tone-free (preCS) period. Safety conditioned rats had higher suppression scores than fear conditioned rats, whereas tone-only controls had scores near zero. Data are presented as mean ± SEM. *p < .05, **p < .01, and ***p < .001; ns, not significant.

To examine the efficacy of the safety cue in modulating the fear state during recall, we calculated a suppression score for each subject. Positive suppression scores would indicate reduced freezing during the tone relative to the preceding preCS period and is consistent with the CS acting as a safety signal, whereas negative suppression scores would indicate increased freezing during the tone and would be consistent with conditioned fear expression. A one-way ANOVA revealed significant difference between the groups in suppression scores [F(2,31)=10.01, p<.0004, η^2^ = .39]. Post-hoc analyses showed that the safety conditioned group had significantly greater suppression scores than the fear group [Student t-test, p<.001, **Fig. 1D**], but not the tone-only groups [p=.303]. In contrast, the fear conditioned group showed negative suppression scores during recall, while the suppression scores for the tone-only group remained close to zero consistent with the absence of associative learning by this group [**Fig. 1D**].

To rule out that the cue-evoked suppression in freezing in safety conditioned animals was not result of some non-associative or unrelated process, we tested a naïve group of rats in which the tone stimulus was presented only at the time of recall. During training, all rats showed the expected increase in post-shock freezing following each footshock indicating successful fear acquisition [**Supplementary Fig. S2A**]. During the recall test, the overall percentage of time spent freezing did not differ significantly for rats tested in the presence or absence of the tone stimulus [No Tone: 67.8% ± 11.8% vs. Tone: 60.9% ± 9.13%, p=.657, n=5 rats per group]. Freezing levels during the tone-free and tone-on periods was not significantly different [Tone: preCS vs. CS, F(1,4)=1.42, p=.299, **Supplementary Fig. S2B**]. These findings demonstrate that presentation of an auditory stimulus is not sufficient to disrupt freezing confirming that fear suppression after safety conditioning is dependent on the previously learned tone-safety association.

### Learned safety cues suppress fear responses to a novel context

Our findings suggest that safety cues can attenuate threat responses which is consistent with the idea they may function as conditioned inhibitors. However, it remained unclear whether this cue could transfer and signal for safety in a new environment associated with threat. To test this possibility, rats previously underwent safety conditioning over three days in chamber A and then were conditioned to fear a novel context (Chamber B) one week later. Safety conditioned rats showed low levels of baseline freezing in Chamber B and displayed robust during conditioning in this new context as indicated by a significant increase in post-shock freezing compared to the baseline period [RM-ANOVA, F(2,36)=15.96, p<.001, **Fig. 2A**].

**Fig. 2.**
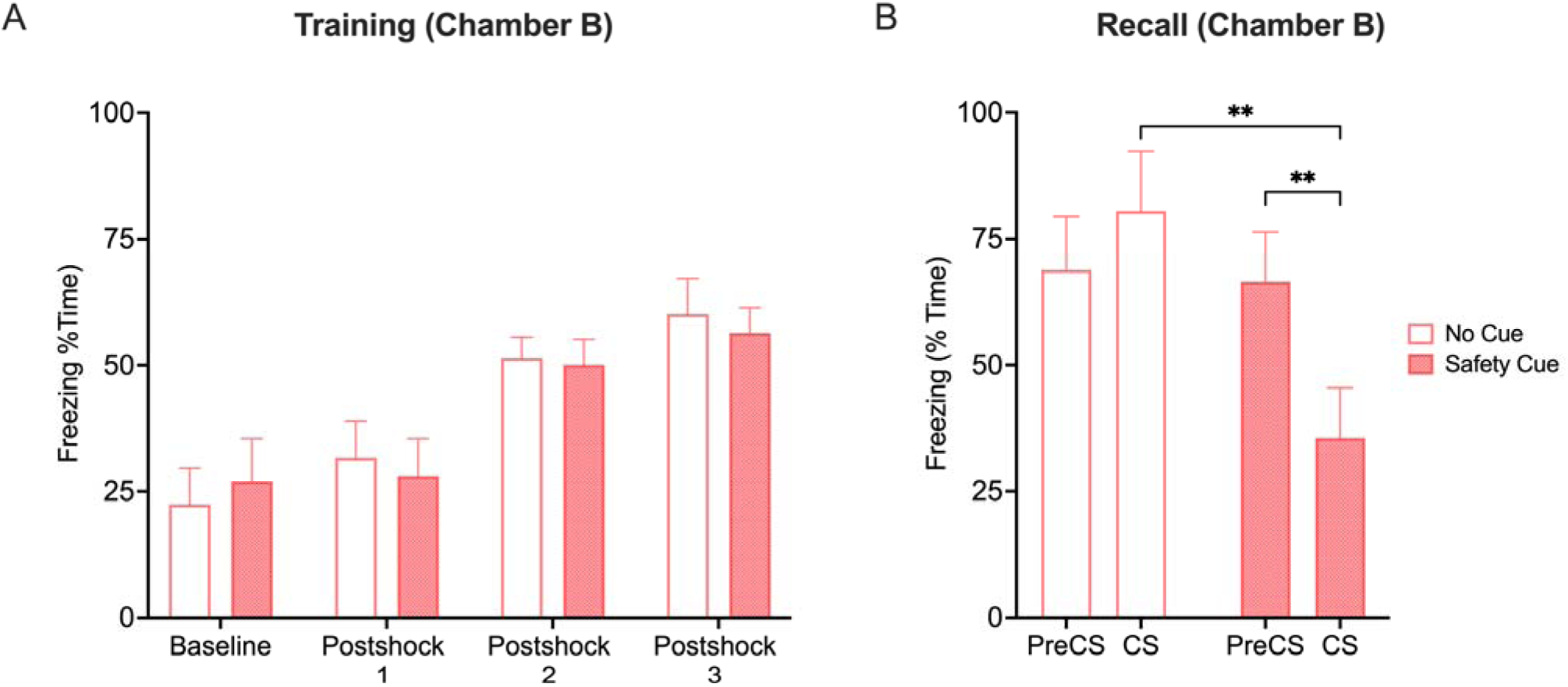
Learned safety cues suppress freezing in a novel threat-associated context. (A) Mean percentage of time spent freezing during conditioning in a novel context (Chamber B). Rats had undergone safety conditioning one week earlier. (B) Mean percentage of time spent freezing during the safety cue and the immediately preceding tone-free period during the recall test conducted 24 h later. Rats tested with the safety cue displayed lower freezing during cue presentation (CS) than during the preceding tone-free (preCS) period. In contrast, rats tested without the cue showed comparable freezing across the equivalent preCS and CS periods. For time-matched intervals, rats tested without the cue displayed greater freezing than rats tested with the safety cue. Data are presented as mean ± SEM. **p < .01.

Twenty-four hours after conditioning in Chamber B, rats were assigned to one of two recall conditions. The tone group was tested in the presence of the previously learned safety cue, whereas the no-tone group was tested without the cue. To determine whether the safety cue reduced contextual freezing, we compared the average percentage of time spent freezing during the three tone presentations with freezing during the intervals immediately preceding each tone. Equivalent time-matched intervals were analyzed for the no-tone group to permit direct comparison of changes in freezing across the recall test.

Before presentation of the safety cue, the tone and no tone groups showed high levels of freezing in Chamber B indicating they had successfully learned to fear the new context [No Tone: 71.5% ± 12.3% vs. Tone: 69.3% ± 9.28%, p=.695]. Analysis of freezing during the preCS and CS intervals revealed a significant Group × Interval interaction [F(1,8)=23.05, p<.005, η² =.74], with no main effect of Interval [F(1,8)=4.72, p=.062] or Group [F(1,8)=2.73, p=.137]. As shown in **Fig. 2B**, rats tested with the safety cue showed reduced freezing during the CS compared to the preceding preCS interval [p<.001]. In contrast, rats tested without the safety cue showed no difference in freezing across the corresponding time-matched intervals [p=.100]. Analysis of suppression scores further confirmed this effect, with rats tested in the presence of the safety cue demonstrating greater suppression than those tested without the cue [Student t-test, t(8)=4.80, p<.001, Cohen’s d=3.04]. These findings indicate that a previously acquired safety cue can suppress freezing in a novel threat-associated context suggesting that the retrieval of the safety memory can modulate fear expression in a context independent manner.

### Molecular changes associated with safety learning

To determine how safety and fear learning impacts the medial prefrontal cortex (mPFC), basolateral amygdala (BLA), and central amygdala (CeA)—key brain regions involved in threat learning—we quantified the immediate early gene product c-Fos expression in fear conditioned (n=8) and safety conditioned (n=11) groups after recall testing. A tone-only group (n=4) was also included in which these rats were presented with the same tones in the training context but never received shock pairings. The purpose of this group was to control for stimulus driven Fos induction that is independent of learning. A one-way multivariate GLM comparing Fos+ counts in the mPFC, BLA, and CeA showed a main effect of Group [Wilks’ λ=.421, F(6, 38)=3.43, p=.008, η_p_²=.351]. Follow-up *post hoc* tests showed that the fear and safety conditioned groups did not differ from each other in Fos+ counts in the mPFC, but they both had significantly higher Fos+ counts than the tone-only control group [fear vs. tone-only, p=.014; safety vs. tone-only, p=.016, respectively, **Fig. 3A**]. Fear conditioned rats had higher Fos+ counts in the BLA and CeA compared to the safety conditioned and tone-only groups [BLA: p=.013 and <.001; CeA: p=.008 and p=.001, respectively, **Fig. 3B**].

**Fig. 3.**
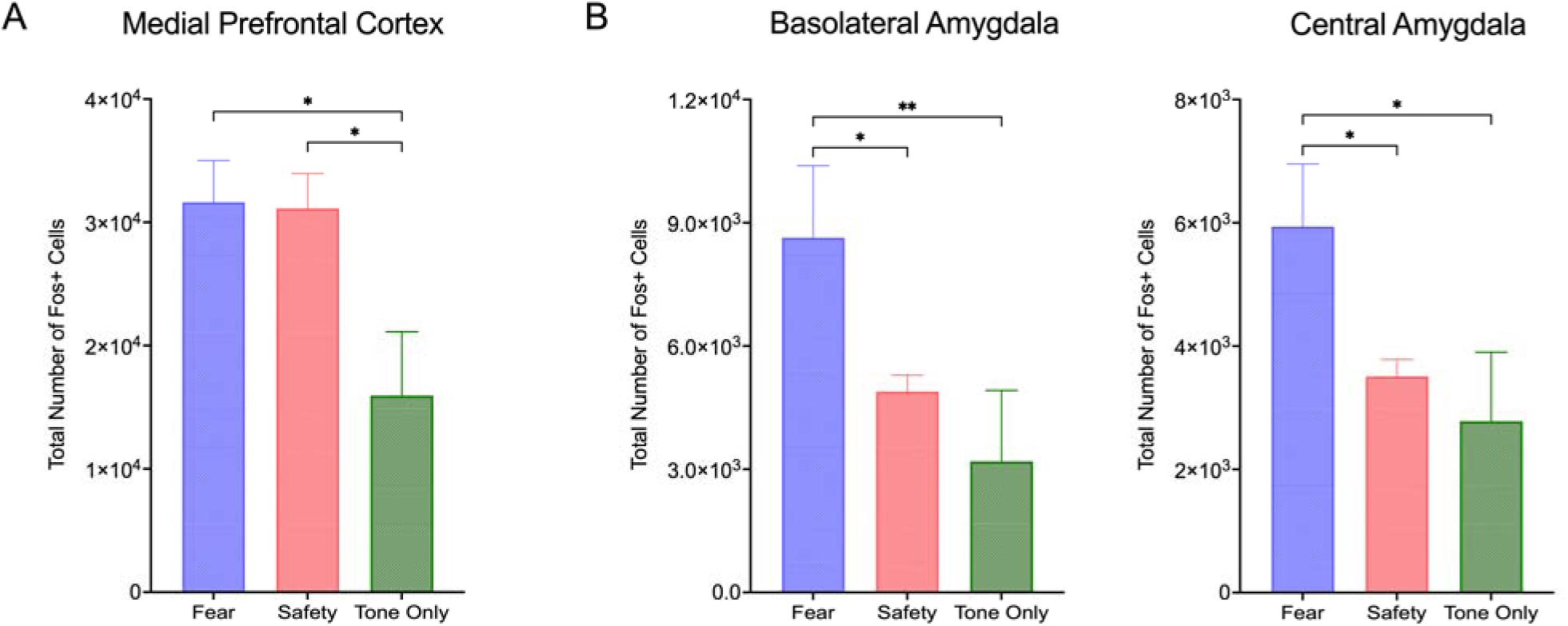
Fear and safety learning produce distinct patterns of Fos activation in medial prefrontal and amygdala regions during recall. (A) Total Fos+ cell counts in the medial prefrontal cortex (mPFC) following recall testing. Both fear and safety conditioned rats had more Fos+ cells than tone-only controls. (B) Total Fos+ cell counts in the basolateral amygdala (BLA) and central amygdala (CeA) following recall testing. Fear conditioned rats had more Fos+ cells in both amygdala subregions than safety conditioned rats and tone-only controls. Data are presented as mean ± SEM. *p < .05 and **p < .01.

Given the importance of AMPA and NMDA receptors in mediating associative plasticity and synaptic changes that support cued fear conditioning, we wondered if safety learning might impact the expression of these glutamatergic receptors. Western blot analysis revealed no group differences in PSD-95 [p=.381], GluN1 [p=.699], or GluA1 [p=.424] expression in mPFC homogenates [**Fig. 4A**], but c-Fos protein levels were increased in both safety and fear conditioned groups compared to the tone-only controls [ps<.043]. In contrast, analysis of amygdala homogenates revealed a significant Group × Protein interaction [F(6,36)=2.94, p=.019, η²p=.329], indicating that group differences varied across the examined proteins. Simple effects analyses revealed group differences for PSD-95 [F(2,36)=7.00, p=.003, η²p=.280], GluN1 [F(2,36)=3.59, p=.038, η²p=.166], GluA1 [F(2,36)=5.01, p=.012, η²p=.218], and c-Fos [F(2,36)=3.92, p=.029, η²p=.179]. As shown in **Fig. 4B**, amygdala PSD-95 expression was increased for the safety conditioned group compared to the fear conditioned [p=.024] and tone-only [p<.001] groups. GluA1 expression was also significantly elevated in safety conditioned rats compared to the tone-only group [p=.002] with a trend toward higher expression than the fear conditioned group [p=.072]. In contrast, GluN1 expression was higher in fear conditioned rats than both safety conditioned rats [p=.025] and tone-only controls [p<.001]. Whole amygdala c-Fos expression was also increased in fear conditioned rats compared to the safety conditioned rats [p=.029] and tone-only controls [p=.015] consistent with the selective increases in BLA and CeA Fos+ cell counts observed previously.

**Fig. 4.**
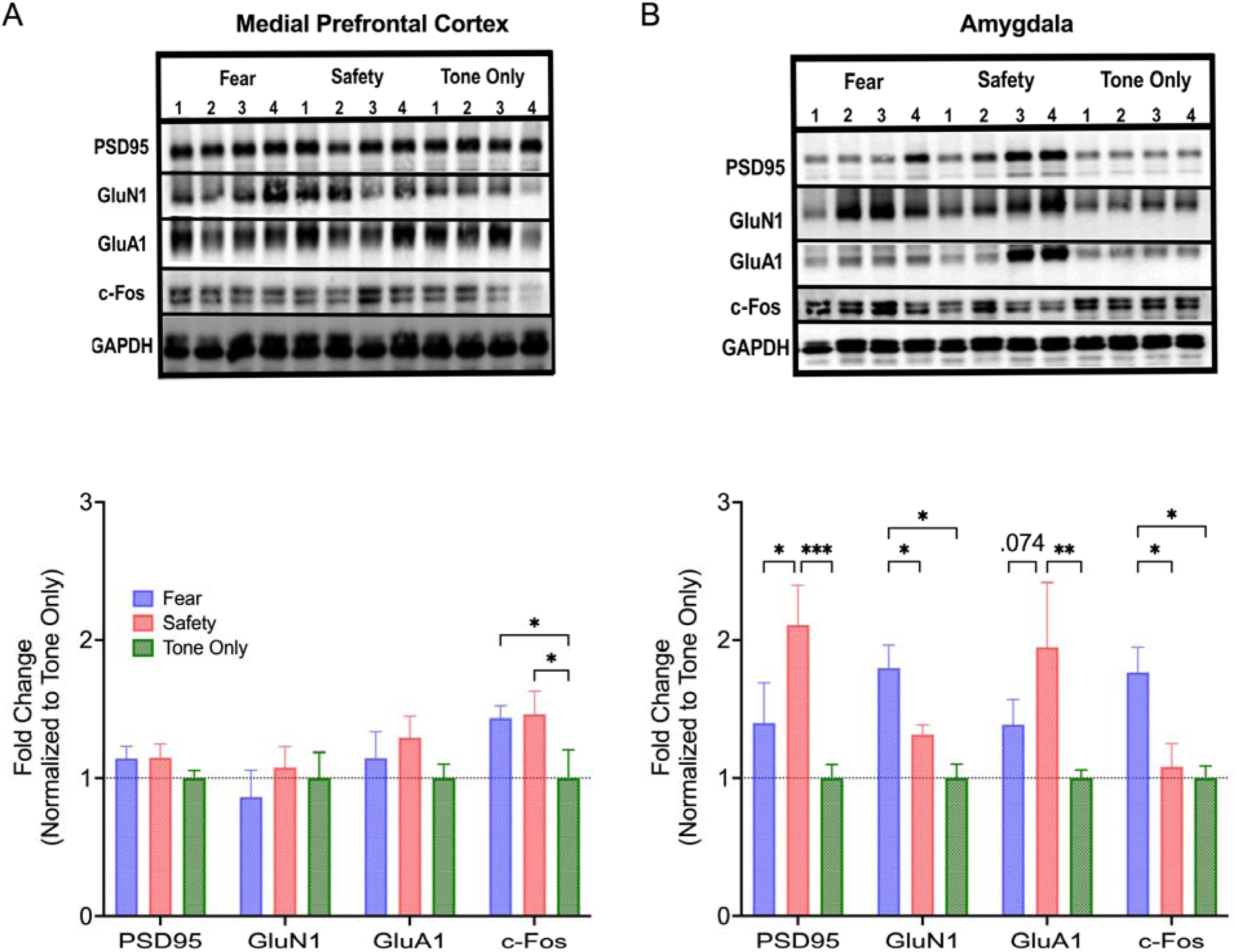
Fear and safety learning produce distinct patterns of glutamatergic postsynaptic protein expression in the medial prefrontal cortex and amygdala during recall. (A) Representative immunoblots and quantification of PSD-95, GluN1, GluA1, and c-Fos in whole mPFC homogenates following recall testing. Both fear and safety conditioned rats showed increased mPFC c-Fos expression relative to tone-only controls. (B) Representative immunoblots and quantification of PSD-95, GluN1, GluA1, and c-Fos in whole-amygdala homogenates. Safety conditioning increased amygdalar PSD-95 and GluA1 expression relative to tone-only controls, whereas fear conditioning increased amygdalar GluN1 and c-Fos expression relative to both safety conditioned rats and tone-only controls. Protein levels were normalized to GAPDH and expressed as fold change relative to tone-only controls. Data are presented as mean ± SEM. *p < .05, **p < .01, and ***p < .001.

### Prior safety learning enhances extinction

Given that presentation of the safety cue rapidly attenuated freezing during recall, we hypothesized that prior safety learning might accelerate extinction as characterized by a more rapid reduction in freezing across repeated cue exposure in the threat-associated context. To test this, safety and fear conditioned rats underwent 5 days of extinction training (12 CS tone presentations per day).

CS-evoked freezing declined markedly across the extinction sessions [Day: F(4,32)=4.09, p=.009, η² =.34] and was overall lower for the safety conditioned group [Group: F(1,8)=7.09, p=.029, η² =.47]. Although the Day × Group interaction was not significant [F(4,32)=1.68, p=.179], a quadratic trend [F(1,8)=7.10, p=.029] was found revealing that safety conditioned rats showed less freezing primarily over the first three extinction sessions than fear conditioned rats [**Fig. 5A**]. Consistent with this pattern, suppression scores also differed between groups on Days 1 to 3 [ps<.046] but not Days 4 to 5 [ps>.32, **Fig. 5B**] of extinction. For the safety conditioned group, daily suppression scores correlated positively with the amount of pre-tone freezing only on Days 2 and 3 of extinction training [Pearson rs=.78-.89, ps<.013] indicating that rats displaying higher levels of context driven freezing during early extinction sessions also showed the largest reduction in freezing during presentation of the safety cue.

**Fig. 5.**
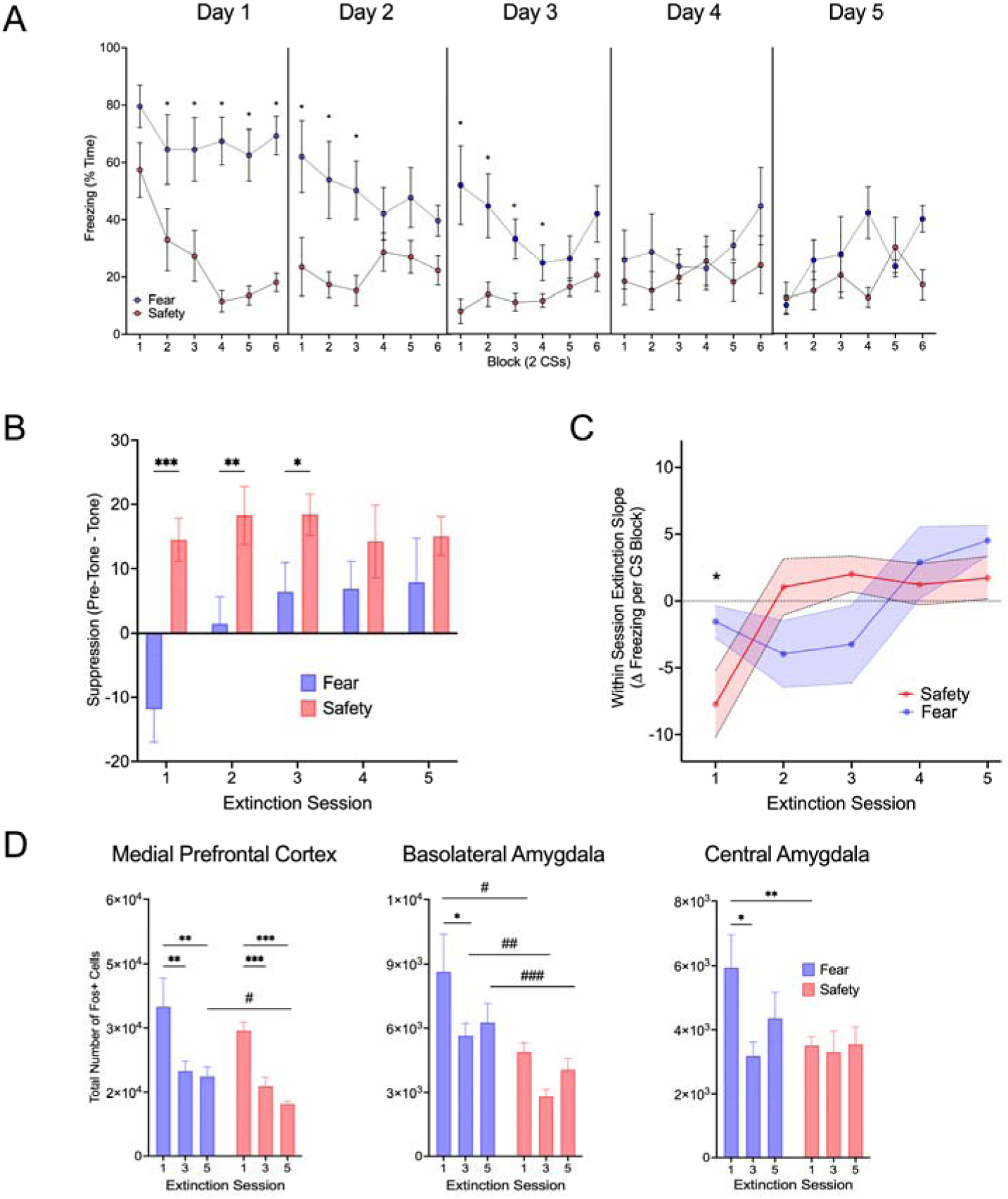
Safety learning accelerates extinction of conditioned freezing and is associated with dissociable patterns of Fos+ cell counts in the medial prefrontal cortex and amygdala. Rats previously trained with explicitly unpaired safety conditioning or standard fear conditioning underwent 5 days of extinction training (12 non-reinforced tone presentations per session, in 6 blocks of 2 trials). (A) Tone-evoked freezing declined across extinction sessions in both groups, but fear conditioned rats showed greater overall freezing. (B) Daily suppression scores (preCS minus CS freezing) were higher in safety than fear conditioned rats on Days 1–3, but not Days 4–5. (C) Within-session extinction slopes (more negative = faster within-session decline). Safety conditioned rats showed steeper slopes on Day 1, plateauing near zero by Day 2, whereas fear conditioned rats declined more gradually across sessions. (D) Total Fos+ cell counts in the mPFC, BLA, and CeA after 1, 3, or 5 days of extinction. Both groups showed higher mPFC Fos on Day 1 than later sessions, with elevated counts persisting longer in fear conditioned rats. Fear conditioned rats also showed higher BLA Fos throughout extinction and higher CeA Fos on Day 1. CeA Fos was stable across extinction session for safety conditioned rats. Data are presented as mean ± SEM. *p < .05, **p < .01, and ***p < .001 for within-group comparisons; #p < .05, ##p < .01, and ###p < .001 for comparisons between the fear and safety conditioned groups.

We also assessed within-session extinction by calculating linear slopes of tone-evoked freezing across each session. In this situation, a negative slope would reflect steeper reductions in tone-evoked freezing which is indicative of faster or more rapid within-session extinction. Analysis of these slopes revealed a significant Day × Group interaction [Wilks’ λ=.179, F(4,5)=5.75, p<.05, η² =.82] along with a main effect of Day [F(4,32)=5.25, p<.002, η² =.40]. The source of the interaction was primarily due to safety conditioned rats extinguishing rapidly during the first extinction session with slopes reaching plateau (near 0) by Day 2, whereas fear conditioned rats displayed more gradual extinction with slopes turning positive only by the fourth extinction session [**Fig. 5C**].

These group differences in extinction learning were also reflected in patterns of Fos immunoreactivity in the mPFC, BLA, and CeA. The results of a multivariate GLM for Fos+ counts across these brain areas revealed significant main effects of Day [Wilks’ λ=.428, F(6,60)=5.28, p<.001, η² =.346] and Group [Group: Wilks’ λ=.757, F(3,30)=3.21, p<.037, η² =.243]. In the mPFC, Fos+ counts were higher on the first extinction session than the third and last sessions for fear and safety conditioned groups [Day 1 vs. Days 3,5 ps<.001]. Fear conditioned rats also showed a nonsignificant trend toward more mPFC Fos+ cells than the safety conditioned rats after the third extinction session [Day 3, p=.079], which reached significance on the last session [Day 5, p=.018]. In the amygdala, differences were most pronounced on the first extinction session with fear conditioned rats having significantly higher Fos+ counts than safety conditioned rats in both the BLA and CeA [ps<.004; **Fig. 5D**]. Group difference in Fos+ counts in the BLA also persisted across the remaining extinction sessions [Day 3: p=.006; Day 5: p=.034, **Fig. 5D**], whereas differences in Fos+ cells between the fear and safety conditioned rats for the CeA were no longer apparent by the third extinction session [p=.135]. This effect was driven primarily by the decline in Fos+ cells in the CeA of fear conditioned rats [Day 1 vs. Day 3: p=.013, **Fig. 5D**] while CeA Fos+ cell counts remained relatively stable across extinction sessions 1, 3, and 5 for the safety conditioned rats [p>.972].

To determine if suppression of conditioned freezing by the tone was associated with different patterns of Fos induction within these brain areas during extinction for safety and fear conditioned rats, Fos+ cell counts were correlated with the change in cue-induced freezing suppression during early (Days 1-3,), late (Days 3-5) and overall (Days 1-5) extinction sessions. For safety conditioned rats, we found that BLA Fos+ counts correlated positively with tone-induced suppression of freezing during early extinction sessions [Day 1-3: Pearson r=.65, p<.05], while mPFC Fos+ counts correlated negatively with tone-induced suppression during the later extinction sessions [Day 3-5, r=-.96, p<.01] and overall tone-induced suppression from the first to last sessions [Day 1-5, r=-.91, p<.05]. In contrast, for fear conditioned rats, Fos+ counts in these brain areas were not correlated with differences in suppression during the tone during early extinction sessions. However, suppression of freezing during the tone was negatively correlated with the number of Fos+ cells in the CeA during later sessions of extinction in fear conditioned animals [Day 3-5, Pearson r=-.88, p<.05].

Together, these findings suggest that safety and fear learning appear to recruit distinct prefrontal-amygdala circuits during extinction. In safety conditioned rats, early suppression of freezing by the tone cue was associated with BLA Fos expression, while later suppression by this tone cue was associated with reduced mPFC activation. In contrast, for fear conditioned animals suppressed freezing during the tone cue was associated with decreased CeA recruitment suggesting dissociable contributions of the prefrontal cortex and amygdala to extinction learning that depends on prior conditioning history.

## 3. Discussion

The present study sought to deepen our understanding of the neural substrates that distinguish safety learning from traditional forms of fear learning and extinction. In this study, we found that safety conditioned rats displayed robust, cue-specific reductions of freezing when exposed to the original conditioning context. This safety cue also possessed the capacity to generalize to a novel threat-associated context and suppress freezing levels when presented. We also found that safety and fear learning resulted in dissociable patterns of neuronal activation and changes in postsynaptic glutamatergic receptors primarily within the amygdala. Finally, safety learning was followed by a more rapid decline in tone-evoked freezing during early extinction training, which was further associated with differences in medial prefrontal and amygdala activation compared to fear conditioned animals. Together, these findings indicate that safety learning is not simply the opposite of fear learning or a byproduct of extinction but represents a qualitatively different learning process which engages specific neural circuits to ultimately impact how conditioned fear is either maintained or expressed.

### Behavioral dissociation between learned fear and learned safety

Safety learning represents a distinct form of associative learning that enables animals to learn that specific cues, contexts, or actions predict the absence of threat. Although various approaches have been used to study safety learning in rodents [28], we used a procedure in which an aversive US (footshock) and a CS (tone) are presented within the same conditioning session but in an explicitly unpaired manner. Under these contingencies, the conditioning context functions as a conditioned excitor (CS+) with the tone signaling for the occurrence of the US-free period. This procedure has been shown to be effective in establishing safety signals in rodents [6,7,9,10,23,29–31]. Consistent with this, we found that safety conditioned rats displayed tone-evoked freezing that was consistently lower than freezing during the corresponding pre-tone (preCS) period, which aligns with the well-established view that stimuli that reliably signal for the absence of an aversive event acquire inhibitory associative strength, rather than simply failing to elicit conditioned fear [32,33]. This stands in marked contrast to fear conditioned rats, in which the arrival of the tone predicts impending danger resulting in higher freezing to the tone and also to the conditioning context. Importantly, group differences in tone-evoked freezing could not be accounted for by overall weaker conditioning of the safety group, as both safety and fear conditioned rats displayed comparable levels of preCS freezing as well as post-shock freezing after shock delivery. Rather, our findings support a dual learning model in which safety conditioned rats do in fact learn that the context is aversive (CS+) while simultaneously learning that the presentation of the tone can come to reliably predict a period free of shock (CS-) enabling the tone to function as a safety signal that actively suppresses freezing and defensive responses whenever it is presented [34].

One possibility is that the reduction in tone-evoked freezing was not the result of safety learning per se but instead came about through non-associative effects on conditioned freezing due to competing orienting and motor responses elicited by the tone itself as recently proposed by Mombelli and colleagues [35]. We believe this is unlikely for two reasons. First, in a separate cohort of rats in which a tone was introduced only during testing but never during conditioning, presentation of this tone had little effect on freezing, i.e., freezing levels during the presentation of the tone did not differ from freezing during the tone-free periods. In fact, overall freezing was similar for rats tested in the presence or absence of the tone. This finding is supported by past evidence that simple CS non-reinforcement trials are insufficient to produce conditioned inhibition, and that the explicit unpaired relationship between the CS and US is necessary for establishing the CS as an active safety signal [36,37]. Second, safety conditioned rats showed a progressive decrease in tone-evoked freezing across repeated presentations with freezing during the preCS period remaining relatively stable both within and across conditioning sessions. If presenting the tone produced external inhibition through eliciting orienting or some other type of distractive response which leads to reduced freezing, then one would expect its suppressive effect on freezing to also habituate with repeated exposure. However, we found the opposite pattern, which is consistent with the idea that tone acquired inhibitory properties rather than lost salience through habituation.

An important feature of a conditioned inhibitor is its capacity to inhibit responses elicited by other independent excitatory conditioned stimuli [33]. Indeed, we found that the safety cue was not only bound to its original training environment but could transfer its inhibitory effect to suppress freezing in a newly conditioned, physically distinct context. This transfer effect on conditioned freezing was absent in rats tested without the safety cue present indicating that prior safety learning does not impair the acquisition of the new contextual fear memory. Instead, these findings suggest that the tone can signal for “safety” whenever it is present regardless of the context. Similar effects have been reported. For example, Takemoto and Song [38] found that mice trained on an auditory fear discrimination paradigm, in which one tone cue (CS+) was paired with a shock and a second tone cue (CS-) was kept explicitly unpaired with the shock, displayed significant suppression of freezing during presentation of the safety cue (CS-) in the training environment as well as in a novel environment. Together, with earlier evidence [10,29] and our finding that the safety cue actively inhibits rather interferes in the capacity to generate fear responses, these results suggest that safety associations, once formed, are not bound to the context in which they are acquired. Instead, learnt safety associations have the capacity to generalize which enables flexible control over fear expression in novel situations.

### Regional differences after safety learning

We found that both fear and safety learning increased Fos+ cell immunoreactivity and whole Fos protein levels in the mPFC compared to tone only controls. In contrast, rats that underwent fear conditioning showed increased Fos immunoreactivity in the BLA and CeA, while safety conditioned rats displayed attenuated Fos expression within these same amygdalar regions. In fact, Fos+ counts for the safety conditioned group were comparable to that of tone-only controls, which we also corroborated through immunoblotting of Fos protein in whole amygdala homogenates. These findings are consistent with previous reports that neuronal activity in the lateral amygdala is reduced following safety learning [5,10,39] and raise the possibility that safety signals might alter the function of neural circuits in the amygdala with threat and defensive behavior.

Our immunoblotting results offer some support to this idea. Although GluA1, GluN1 and PSD-95 levels were unchanged in prefrontal homogenates after safety or fear conditioning, their expression was altered within amygdala. Fear conditioning increased expression of the obligatory NMDA receptor subunit GluN1 in amygdala homogenates compared to safety conditioned and tone-only control animals—a finding consistent with the large body of evidence implicating NMDA receptor dependent long-term potentiation within the amygdala as a cellular mechanism underlying the acquisition and consolidation of conditioned fear memories [40–43]. In contrast, safety conditioning selectively increased amygdalar expression of GluA1 and the postsynaptic scaffolding protein PSD-95, both of which have been associated with AMPA receptor trafficking and the stabilization of glutamate receptors and other signaling molecules at the postsynaptic density [44–47]. The increase in expression of these synaptic markers suggests that at some level safety learning requires active synaptic plasticity and/or reorganization within specific amygdala microcircuits in order to preserve the significance of the safety cue and inhibit fear responding.

### Safety and fear learning differentially recruit the mPFC and amygdala during extinction

Much like safety learning, fear extinction also produces inhibition of conditioned fear responses. However, extinction occurs following prior fear learning and involves repeated non-reinforced presentations of the CS, during which the animal must learn that a previous threat-associated cue no longer predicts an aversive outcome. In contrast, safety learning involves prior experience in which the CS never coincided with the US enabling the animal to learn that the CS always predicted a period free of threat and hence does not require revision of learned meaning of the CS.

Given the different learning contingencies involved in these approaches, we wondered whether the dynamics of extinction might be affected for rats that had previously undergone safety or fear conditioning. Safety conditioning resulted in an immediate and robust suppression of tone-evoked freezing both within and across the first three sessions of extinction training, whereas extinction training of fear conditioned animals required at least four sessions before freezing to tone declined to the levels comparable to those observed in the safety group. In fear conditioned rats, BLA and CeA Fos+ counts declined across extinction sessions, although only CeA Fos+ cell counts reached values comparable to the safety group. Greater fear suppression to the tone appeared to be associated with lower Fos+ cell counts in the CeA for fear conditioned animals, but only during the later stages of extinction learning. This is consistent with evidence that successful fear extinction depends on a gradual attenuation of amygdala activity, particularly within the CeA, which promotes error-driven updating of the original threat memory as the animal learns that previously shock-predictive cues no longer signal for threat [22,48–51].

Safety conditioned rats, by contrast, showed overall low BLA and CeA Fos expression across all examined extinction sessions which appears to parallel the accelerated extinction to both the tone and context observed in this group. The BLA is important for evaluating the degree of threat associated with specific environmental cues and/or situations [52–55]. However, emerging evidence also indicates that the BLA contains distinct neuronal populations that also encode for the rewarding or safety features of stimuli [56–59]. Interestingly, some BLA “safety” neurons appear to shift their response from threat-predictive cue during extinction while others remain persistently selective for the “safety” cue only [60]. This suggests that rather than requiring the slower, error-driven updating process thought to support traditional (CS-no US) extinction [61], safety learning may leverage these stable, preexisting safety ensembles within the BLA to rapidly suppress freezing. This interpretation fits with our finding that fear suppression during early extinction sessions was positively correlated with BLA Fos expression in safety conditioned rats likely reflecting the recruitment of theses safety encoding neurons rather than erosion of the threat memory.

Accumulating evidence has also pointed to reciprocal connections between the mPFC and amygdala in supporting the retrieval of extinction memory [21,55,62]. We found that later extinction sessions were associated with reduced mPFC Fos expression in safety-conditioned rats, whereas fear-conditioned rats continued to exhibit comparatively elevated mPFC Fos expression even after the final extinction session. This pattern is consistent with findings from animal and human studies showing that omission of an expected aversive event recruits prefrontal circuits to suppress fear elicited by a previously threat-predictive cue [63–68]. Because the tone had already been established as a reliable predictor of non-threat in the safety-conditioned group, its repeated presentation during extinction would be expected to generate little prediction error, thereby reducing the demand placed on prefrontal circuits. In contrast, resolving the discrepancy between the tone’s previous association with threat and its new non-reinforced outcome may require greater or more sustained engagement of prefrontal and amygdala circuits in fear-conditioned animals.

The eventual suppression of freezing in fear conditioned rats should not necessarily be interpreted as evidence that extinction transformed the tone into a safety cue equivalent to that established through safety conditioning. Extinction is generally understood to produce a new CS–no US association that competes with, rather than erases, the original threat association. In addition, retrieval of extinction memory is also highly context dependent with conditioned fear often recovering when an extinguished cue is presented outside the extinction context, including when it is presented in a novel context [65,69,70]. Thus, an extinguished fear cue may suppress responding within the extinction context without acquiring the more transferable inhibitory properties of an explicitly trained safety cue. This distinction is important given that we showed that a learned safety cue was capable of generalizing and reducing conditioned freezing to a novel context. This raises the intriguing possibility that prior safety learning may effectively prime extinction-related processes by recruiting established safety-encoding ensembles and modulating downstream amygdala circuits, thereby permitting more rapid and potentially more generalizable suppression of threat responses.

### Implications and Conclusions

Impaired discrimination of safe or previously safety associated cues is believed to contribute to the excessive fear that is a hallmark of anxiety- and trauma-related disorders, such as phobias, panic disorder, and PTSD [6,23]. Our finding that fear conditioned, but not safety conditioned, rats showed elevated BLA and CeA Fos expression parallels neuroimaging evidence of exaggerated amygdala reactivity and in patients suffering from PTSD [71]. This suggests that effective safety learning may depend on the ability to disengage amygdala-mediated threat responses when cues signal the absence of danger.

These findings suggest that treatments for pathological fear could benefit from developing strategies that directly target safety learning rather than relying only on traditional extinction-based methods [8,72,73], which remain highly context dependent and vulnerable to relapse [61,74,75]. Safety learning reflects the acquisition of information of a stimulus’s protective properties rather than the de-valuation of a previous threat-associated cue. Because of this and the potential for a safety cue to generalize across contexts and accelerate subsequent extinction learning, interventions designed to strengthen the acquisition, retrieval, and generalization of safety memories could offer a complementary and potentially more durable approach to reduce pathological fear [7,76,77].

## 4. Materials and Methods

### Subjects

Sprague Dawley rats, weighing approximately 200-250 g, were obtained from Charles River Laboratories (QC, Canada). All subjects were group-housed (2 rats per cage) with *ad libitum* access to food and water and were maintained in a temperature-controlled room (25 ± 2^°^C) at 60% humidity. Behavioral testing began after two weeks of handling. Experiments were performed between 0900 and 1800 h during the photophase of a 12-hour light/dark schedule (lights on at 0700 h). All experimental procedures were approved by the Trent University Animal Care Committee and were compliant with the Canadian Council on Animal Care guidelines.

### Apparatus

For safety and fear conditioning, rats were individually placed into a Plexiglas operant chamber (designated Chamber A, dimensions: 25.4 x 25.4 x 36.5 cm) housed in a sound attenuating cabinet. Each chamber had a standard grid floor consisting of 21 stainless steel rods (4 mm diameter, 1 cm distance) connected to an adjustable shock generator (Ugo Basile, Varese, Italy) for delivery of a mild scrambled foot shock. A ventilation fan provided a constant background noise of 55 dB in the cabinets, and the chamber was illuminated by a 2.5 W white LED light. A tone was presented through a loudspeaker mounted on the ceiling of the cabinet. Training and testing sessions were recorded by a digital camera placed above the conditioning chamber and connected to a laptop computer. The conditioning chambers were cleaned with Oxivir Five 16 concentrate (Diversey Inc. Canada) after each animal.

In some experiments, rats were tested in an alternate context (designated Chamber B), which had the same physical dimensions as above but included several distinct visual and olfactory modifications. The chambers were further modified to include lined black-and-white checkered walls and the ambient lighting reduced by 60%. The chamber walls and floor were wiped with a 5% lemon scent solution to serve as a distinctive olfactory cue. The chambers were cleaned with 70% ethanol and the lemon scent reapplied between animals.

### Fear and Safety Conditioning

We adapted our procedures after Pollak et al. (2010). Rats were first habituated to the conditioning chamber over three days with each session lasting 22 minutes in duration. Following the last habituation session, rats were randomly assigned to either fear, safety, or control groups. Conditioning occurred over three consecutive days. On each conditioning day, rats assigned to the fear conditioning group received 5 tones (20 s, 5 kHz, 80 dB) that co-terminated with a footshock (2 s, 0.70 mA) on each conditioning day. The mean inter-tone interval was 120 s (range: 90-180 s). For the safety learning group, rats received 5 tones (20 s, 5 kHz, 80 dB) and 5 footshocks (2 s, 0.70 mA) on each conditioning day. Tones and shocks were always presented in an explicitly unpaired manner, such that the tone never predicted the shock and shock delivery never occurred during the onset or duration of the 20 s tone. The mean interval between the footshock and the nearest tone was 120 s (range 100-140 s). The precise timing of the tones and shocks delivered was pseudorandomized across each training day to avoid incidental learning. For some experiments, a third group of rats served as tone-only controls. These rats received the same number of tones as the fear and safety groups but no footshocks were delivered during these sessions. The duration of time spent in the chamber was also matched across groups.

Twenty-four hours after the final conditioning session, a recall test was performed. Rats were returned to the same conditioning chambers. After 90 s elapsed, the rats were presented with three test tones (20 s duration) each separated by an interval of 60 s. No shocks were delivered during the recall test. In some experiments rats underwent extinction training that was carried out over three or five consecutive days. For extinction sessions, all rats received 12 presentations of the tone (20 s duration) each separated by 60 s. No footshocks were delivered during extinction sessions. The 12 tones were divided into 6 blocks of 2 averaged tone trials each for each day of extinction. Extinction training was carried out in the same chambers as used during conditioning.

To examine the transfer effect of a learned safety cue, rats underwent fear conditioning in a distinct chamber then that used for safety learning. Rats were acclimated to this novel chamber (Chamber B) for 180 s before receiving three unsignaled footshocks (0.7 mA, 1 s duration, 60 s inter-shock interval). The rats remained in the chamber for an additional 60 s after the late footshock before being returned to their home cages. Twenty-four hours later, the rats were re-exposed to Chamber B for a recall test. During the recall test, one group received three presentations of a previously learned safety cue (20 s, 5 kHz, 80 dB), whereas the other group was tested without the cue present. Both groups had previously undergone safety learning one week earlier. For rats presented with the safety cue, the first tone was presented 90 s after the animal was placed into the chamber, with subsequent tones presented every 60 s. Rats tested without the safety cue were placed into the chamber for the same duration of time as rats in the safety cue present condition.

To examine interference effects from novel auditory stimulus during testing, each rat was placed into a conditioning chamber and allowed to explore this chamber for 180 s before receiving three unsignaled footshocks (0.7 mA, 1 s, inter-shock interval 60 s). Sixty seconds after the last footshock, the rat was returned to its homecage. Twenty-four hour later, all rats were returned to the original conditioning chamber. One group received three presentations of a novel auditory tone stimulus (20 s, 5 kHz, 80 dB) with the first tone play 90 s after placement in the chamber and subsequent tones presented every 60 s thereafter. The other group was not exposed to the tone but remained in the chamber for the same duration.

Defensive freezing—defined as the absence of observable movement except those necessary for respiration—was measured using an automated freeze detection system (Any Maze, Stoelting, Wood Dale, IL, USA). Animals were scored as “freezing” when the movement index fell below a threshold of 70 arbitrary units for a minimum duration of 1 s. The amount of time freezing to each tone was expressed as a percentage of freezing to the tone. In addition to tone-evoked freezing, we also measure freezing during the 30 s periods immediately preceding (pre-tone) each tone.

### Fos Immunohistochemistry and Quantification

Ninety minutes after recall testing, rats were anesthetized with of sodium pentobarbital (340 mg/ml, Euthanyl, Merck Animal Health Canada) and underwent transcardiac perfusion with 0.1 M phosphate buffered saline (PBS, pH=7.4) followed by ice cold 4% (w/v) formaldehyde fixative (pH=7.4) freshly prepared from depolymerized paraformaldehyde. Brains were removed and postfixed in the same solution overnight at 4°C and then transferred into 30% (w/v) sucrose and 0.01% (w/v) sodium azide dissolved in PBS. A sliding microtome was used to cut 50 μm thick coronal sections. All sections were stored at −20°C in a cryoprotectant solution containing 30% sucrose, 1% polyvinylpyrrolidone, and 30% ethylene glycol in 0.2 M PBS until processed.

As described elsewhere [78], immunohistochemical staining for c-Fos was performed on free floating sections incubated with a polyclonal rabbit anti-c-Fos antibody (1:15000, EMD Millipore Canada). The primary antibody was diluted in 5% (v/v) normal goat serum, 1% bovine serum albumin and 0.3% (v/v) Triton X-100 dissolved in PBS, and sections were incubated for 48 hrs in this mixture 4°C. Sections were then incubated in biotinylated goat anti-rabbit secondary antibody (1:500, Vector Laboratories) solution for 2 hrs at room temperature, followed by incubation in avidin-biotin complex at for 1 hr (1:500, Vectastain ABC Elicit, Vector Laboratories, Newark, USA). Finally, the reaction was visualized using 2.5% (w/v), 0.02% (w/v) DAB, and 0.000083% (v/v) H2O_2_ in 0.175 M sodium acetate to produce a blue and black product. Sections were mounted on slides, dehydrated through a series of alcohols, cleared in xylene, and coverslipped. Sections from all groups within an experiment were processed at the same time using the same conditions to minimize variability.

Design-based unbiased stereological procedures were performed to determine the number of c-Fos (Fos+) cells using the optical fractionator method [79]. This quantification was done on a Nikon Eclipse 80i microscope equipped with a motorized 3-axis stage, a high-resolution camera, and computerized stereology software (Stereologer, SRC Biosciences, FL USA) by experimenters who were blind to the group condition of the subjects. The reference spaces were defined in accordance with the rat brain atlas based on specific landmarks to identify these areas [80]. The anterior– posterior (AP) level from the bregma of the analyzed regions was as follows: basolateral amygdala (BLA: AP -2.8 mm to -3.60 mm), central amygdala (CeA: AP -2.8 mm to -3.8 mm), and medial prefrontal cortex (mPFC: AP 3.72 mm to 2.52 mm). Regions were traced at low power magnification and Fos cells were counted using an optical disector probe at 100× oil-immersion magnification. For the mPFC, every sixth section was sampled using a 400 µm sampling grid and a optical disector height set to 18 µm. For the BLA and CeA, every twelfth section was sampled using a 150 µm sampling grid and a optical disector height set to 12 µm. Section thickness was measured at each sampling site and used to calculate the tissue sampling fraction for each subject. The total number of Fos cells within each region was estimated using the optical fractionator formula: N_Total_ = ΣQ × 1/ssf × 1/asf × 1/tsf, where ΣQ is the number of Fos cells counted, ssf is the section sampling fraction, asf is the area sampling fraction, and tsf is the tissue sampling fraction. The coefficient of error was calculated and all values were <0.12.

### Protein Isolation and Western Blots

Rats were sacrificed by rapid decapitation 60 min after the recall test. Brains were rapidly removed, and the medial prefrontal cortex (mPFC) and amygdala were dissected on ice, snap-frozen in liquid nitrogen, and stored at −80°C. Tissue was lysed in radioimmunoprecipitation assay (RIPA) buffer containing 50 mM Tris (pH 7.4), 150 mM NaCl, 5 mM EDTA, 1% Triton X-100, 0.1% sodium dodecyl sulfate, 0.5% deoxycholate, and a protease/phosphatase inhibitor cocktail (5872S, Cell Signaling Technology, Danvers, MA, USA). Tissue homogenates were sonicated and centrifuged, and total protein concentrations were determined using a bicinchoninic acid assay (Pierce BCA Protein Assay Kit, Pierce Biotechnology). Samples were subsequently solubilized in 4× Laemmli sample buffer (1610747, Bio-Rad, Hercules, CA, USA).

Equal amounts of protein (40 µg per sample) from the safety- and fear-conditioned groups were separated by 10% SDS-PAGE and electrophoretically transferred onto Immun-Blot PVDF membranes (1620177, Bio-Rad). Membranes were blocked in 5%non-fat dry milk prepared in Tris-buffered saline containing Tween 20 (TBST) for 1.5 hrs and incubated overnight at 4°C with rabbit monoclonal anti-GAPDH (1:20,000, ab181602, Abcam, Cambridge, UK), mouse monoclonal anti-PSD-95 (1:1000, 75-028, NeuroMab, Davis, CA, USA), rabbit monoclonal anti-GluA1 (1:1000, ab109450, Abcam), mouse monoclonal anti-GluN1 (1:1000, MAB1586, Millipore, Burlington, MA, USA), or rabbit polyclonal anti-c-Fos (1:1000, ABE457, Millipore). Following primary-antibody incubation, membranes were washed in TBST and incubated with the appropriate HRP-conjugated goat anti-mouse IgG (W4021, Promega, Madison, WI, USA) or goat anti-rabbit IgG (W4011, Promega, Madison, WI, USA) secondary antibody (1:10000) for 1 hr at room temperature. Immunoreactive bands were visualized using SuperSignal West Femto Maximum Sensitivity Substrate (34095, Thermo Fisher Scientific, Waltham, MA, USA), imaged using an ImageQuant LAS 500 system (GE Healthcare Life Sciences, Pittsburgh, PA, USA), and quantified using ImageJ software (National Institutes of Health, Bethesda, MD, USA).

Band density was quantified using Fiji/ImageJ. Images were converted to 8-bit grayscale and identical rectangular regions of interest were placed over each band corresponding to the expected molecular weight of the target protein. Whenever a closely migrating doublet was present, densitometry was restricted to the band corresponding to the expected molecular weight of the target protein and adjacent nonspecific bands were excluded. Local background was measured for each lane using an identically sized unstained region adjacent to the band. Background corrected density was calculated as the local background raw integrated density minus the band raw integrated density. Target protein values were then normalized to the corresponding housekeeping protein signal (GAPDH) to minimize intra- and inter-gel variability. Data are presented as fold change relative to tone-only control levels.

### Statistical Analyses

All analyses were performed using IBM SPSS Statistics (version 29.0.2.0, IBM Corp., Armonk, NY, USA) or GraphPad Prism (version 10.6.1 for Mac OS, GraphPad Software, Boston, MA, USA) software. Data were initially assessed for normality and homogeneity of variance using Shapiro– Wilk and Levene’s tests. Unless otherwise noted, all data met assumptions for parametric testing.

To examine freezing during conditioning, the average percentage of time spent freezing to the tones was analyzed using a two-way repeated-measures analysis of variance (ANOVA) with Group (fear, safety, tone-only) as the between-subjects factor and Day (1-3) as the within-subjects factor. To further examine for changes in freezing across individual tone presentations, trial-by-trial freezing to each tone was analyzed using a three-way mixed ANOVA with Group as the between-subjects factor and Day (1-3) and Trial (tone presentations 1-5 per day) as within-subjects factors. Post-shock freezing was analyzed separately using a two-way repeated-measures ANOVA with Group (fear conditioned, safety conditioned) as the between-subjects factor and Day (1-3) as the within-subjects factor. Tone-only animals were not included in the post-shock analysis because they did not receive footshock. Where appropriate, significant main effects or interactions were followed by post hoc comparisons that included paired t-tests or one-way ANOVAs.

For recall tests, freezing during the preceding 20 s pre-tone (preCS) and the 20 s tone (CS) periods were analyzed using a mixed design ANOVA with Group as the between subject and Interval (preCS vs. CS) and Block (tone 1-3) as within-subject factors. A suppression score was calculated by subtracting freezing during presentation of the CS from freezing during immediately prior to the CS (preCS-CS). These difference scores were then averaged across all tone trials during the recall session to provide a total suppression value for each subject. Positive values would indicate reduced freezing during the tone (CS) relative to the pre-tone (preCS) period, whereas negative values would indicate increasing freezing to the tone (CS) compared to the pre-tone (preCS) period. Suppression scores were then compared using one-way ANOVA or independent samples t-tests, where appropriate.

Conditioned freezing in the novel context (chamber B) was analyzed using mixed-design ANOVA with Group (tone vs. no tone) as a between subject factor and Interval (preCS vs. CS or equivalent time-matched intervals) as the within-subject factor. Suppression scores were calculated as above and compared using either paired or independent sample t-tests.

For extinction, tone-evoked freezing was analyzed using a mixed-design ANOVA with Group (fear vs. safety) as the between-subject factor and Extinction Day (1–5) as the within-subject factor. Linear within-session extinction slopes were computed for each subject by fitting regression lines across the tone trials for each extinction day. Slopes and suppression scores were analyzed separately using a mixed-design ANOVA with Group as the between-subject factor and Day as the within-subject factor. Correlations were computed between suppression scores and pre-tone freezing to assess whether the magnitude of safety cue-induced freezing suppression varied as a function of context fear.

Total c-Fos (Fos+) cell counts for each brain region was analyzed using a one-way multivariate General Linear Model (GLM) with Group (fear, safety, tone-only) as the between subject factor or as a two-way multivariate GLM with Group (fear, safety) and Extinction session (1, 3, or 5 days) as the between-subjects factors with both analyses followed by Bonferroni-corrected post hoc comparisons, where appropriate. Significant multivariate effects were followed by univariate analyses and simple effects comparisons to determine the direction of brain region-specific effects. Correlation analyses (Pearson’s r) were conducted between early (Days 1-3), late (Days (3-5) and overall (Days 1-5) extinction-related suppression scores and Fos+ counts within each brain region. Western blots for each brain region were analyzed separately using two-way mixed design ANOVAs with Group (safety, fear, tone-only) as the between subject factor and Synaptic Marker (PSD-95, GluA1, GluN1, Fos) as the within-subject factor. Significant results were followed up through simple effects analyses and Bonferroni-corrected post hoc tests. Effect sizes are reported as partial eta-squared (η²p) for ANOVAs and Cohen’s *d* for *t*-tests, where appropriate. Significance thresholds were set at p < 0.05 (two-tailed). Data are presented as mean ± SEM.

## Conflict of Interest

The authors declare no competing interests.

## Acknowledgements

This work was supported by a Discovery Development Grant (DDG-2024- 00012) from the Natural Sciences and Engineering Research Council of Canada (NSERC) and by the John R. Evans Leaders Fund from the Canada Foundation for Innovation (35764) to NMF.

## Author Contributions

NMF conceived the project and designed the research. MA, CC, TAM, and SL performed research and analyzed the data. LJM, HL, and NMF contributed resources. NMF, LJM, and HL wrote and edited the manuscript.

## Data Availability

The data that support the findings of this study are available from the corresponding author upon reasonable request.

---

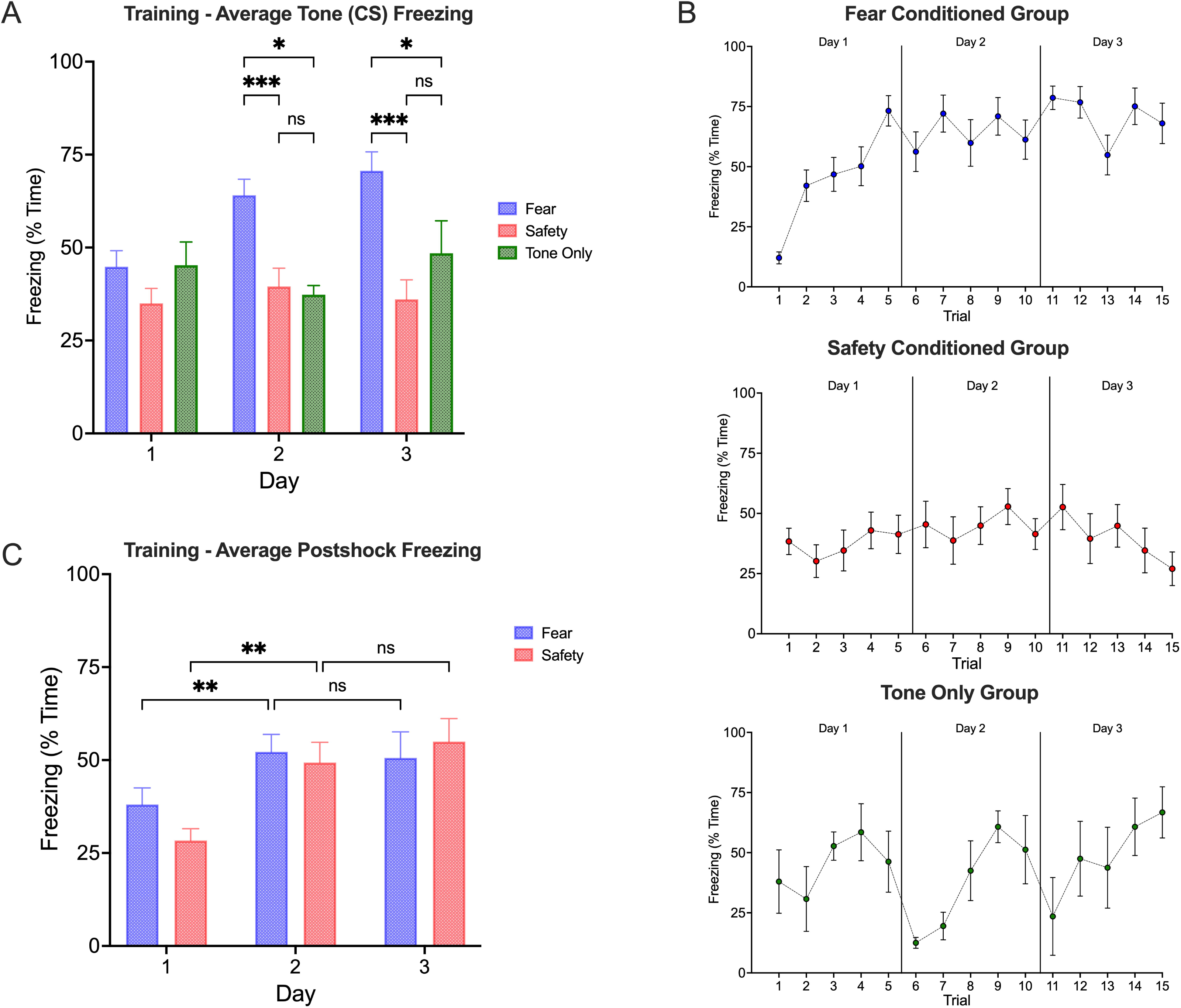

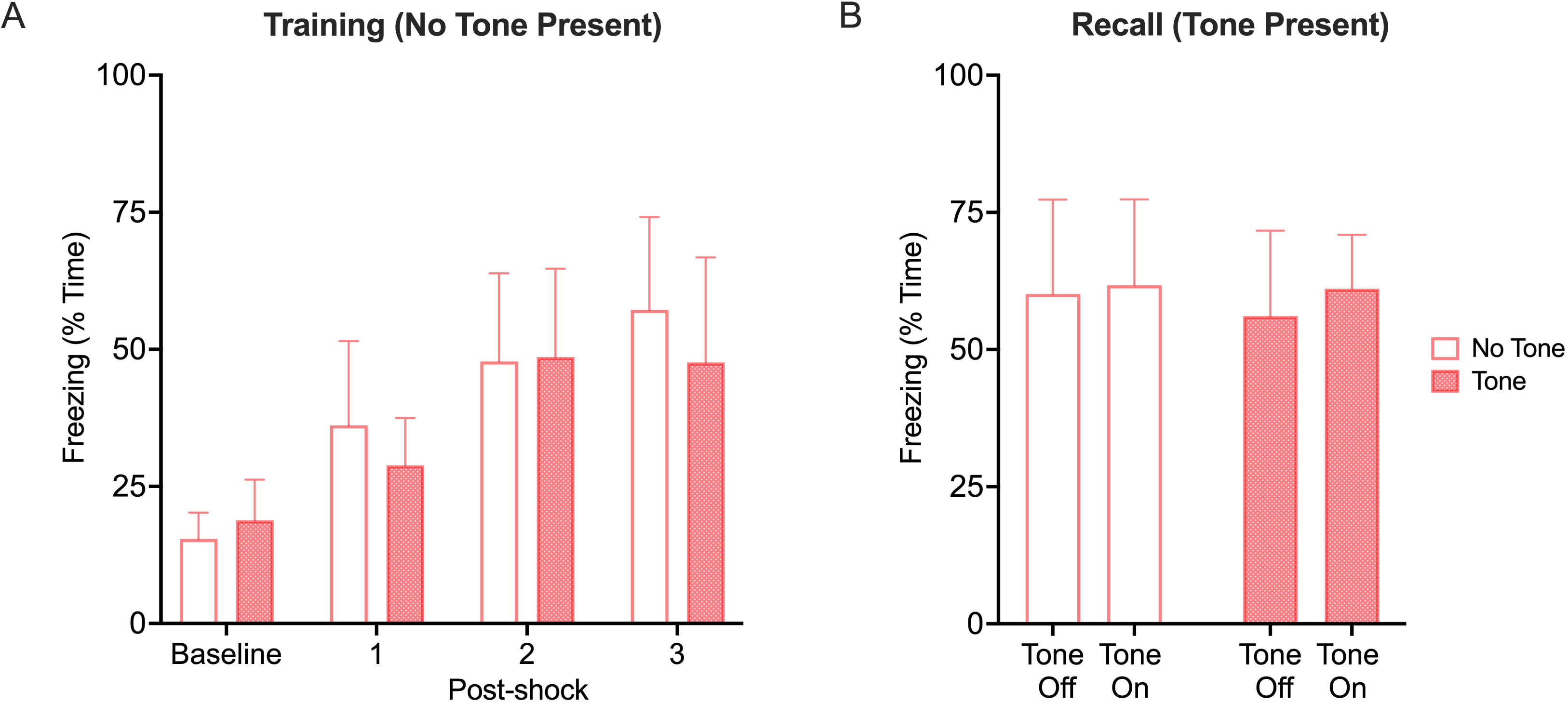

## Notes

### Competing Interest Statement

The authors have declared no competing interest.

## References

1 Fanselow, M. S. & Wassum, K. M. The Origins and Organization of Vertebrate Pavlovian Conditioning. Cold Spring Harb Perspect Biol 8, a021717, doi:10.1101/cshperspect.a021717 (2015).

2 Mobbs, D., Hagan, C. C., Dalgleish, T., Silston, B. & Prevost, C. The ecology of human fear: survival optimization and the nervous system. Front Neurosci 9, 55, doi:10.3389/fnins.2015.00055 (2015).

3 Sangha, S., Diehl, M. M., Bergstrom, H. C. & Drew, M. R. Know safety, no fear. Neurosci Biobehav Rev 108, 218–230, doi:10.1016/j.neubiorev.2019.11.006 (2020).

4 Laing, P. A. F., Felmingham, K. L., Davey, C. G. & Harrison, B. J. The neurobiology of Pavlovian safety learning: Towards an acquisition-expression framework. Neurosci Biobehav Rev 142, 104882, doi:10.1016/j.neubiorev.2022.104882 (2022).

5 Foilb, A. R., Sansaricq, G. N., Zona, E. E., Fernando, K. & Christianson, J. P. Neural correlates of safety learning. Behav Brain Res 396, 112884, doi:10.1016/j.bbr.2020.112884 (2021).

6 Fendt, M., Kreutzmann, J. C. & Jovanovic, T. Learning safety to reduce fear: Recent insights and potential implications. Behav Brain Res 411, 113402, doi:10.1016/j.bbr.2021.113402 (2021).

7 Pollak, D. D. et al. A translational bridge between mouse and human models of learned safety. Ann Med 42, 115–122, doi:10.3109/07853890903583666 (2010).

8 Schiller, D., Levy, I., Niv, Y., LeDoux, J. E. & Phelps, E. A. From fear to safety and back: reversal of fear in the human brain. J Neurosci 28, 11517–11525, doi:10.1523/JNEUROSCI.2265-08.2008 (2008).

9 Christianson, J. P. et al. Inhibition of fear by learned safety signals: a mini-symposium review. J Neurosci 32, 14118–14124, doi:10.1523/JNEUROSCI.3340-12.2012 (2012).

10 Rogan, M. T., Leon, K. S., Perez, D. L. & Kandel, E. R. Distinct neural signatures for safety and danger in the amygdala and striatum of the mouse. Neuron 46, 309–320, doi:10.1016/j.neuron.2005.02.017 (2005).

11 Asok, A., Kandel, E. R. & Rayman, J. B. The Neurobiology of Fear Generalization. Front Behav Neurosci 12, 329, doi:10.3389/fnbeh.2018.00329 (2018).

12 Maren, S. Putting the brakes on fear. Neuron 80, 837–838, doi:10.1016/j.neuron.2013.11.008 (2013).

13 Likhtik, E. & Paz, R. Amygdala-prefrontal interactions in (mal)adaptive learning. Trends Neurosci 38, 158–166, doi:10.1016/j.tins.2014.12.007 (2015).

14 Sengupta, A. et al. Basolateral Amygdala Neurons Maintain Aversive Emotional Salience. J Neurosci 38, 3001–3012, doi:10.1523/JNEUROSCI.2460-17.2017 (2018).

15 Sepahvand, T., Power, K. D., Qin, T. & Yuan, Q. The Basolateral Amygdala: The Core of a Network for Threat Conditioning, Extinction, and Second-Order Threat Conditioning. Biology (Basel) 12, doi:10.3390/biology12101274 (2023).

16 LeDoux, J. E., Iwata, J., Cicchetti, P. & Reis, D. J. Different projections of the central amygdaloid nucleus mediate autonomic and behavioral correlates of conditioned fear. J Neurosci 8, 2517–2529, doi:10.1523/JNEUROSCI.08-07-02517.1988 (1988).

17 Ciocchi, S. et al. Encoding of conditioned fear in central amygdala inhibitory circuits. Nature 468, 277–282, doi:10.1038/nature09559 (2010).

18 Odriozola, P. & Gee, D. G. Learning About Safety: Conditioned Inhibition as a Novel Approach to Fear Reduction Targeting the Developing Brain. Am J Psychiatry 178, 136–155, doi:10.1176/appi.ajp.2020.20020232 (2021).

19 Sotres-Bayon, F. & Quirk, G. J. Prefrontal control of fear: more than just extinction. Curr Opin Neurobiol 20, 231–235, doi:10.1016/j.conb.2010.02.005 (2010).

20 Arruda-Carvalho, M. & Clem, R. L. Prefrontal-amygdala fear networks come into focus. Front Syst Neurosci 9, 145, doi:10.3389/fnsys.2015.00145 (2015).

21 Giustino, T. F. & Maren, S. The Role of the Medial Prefrontal Cortex in the Conditioning and Extinction of Fear. Front Behav Neurosci 9, 298, doi:10.3389/fnbeh.2015.00298 (2015).

22 Marek, R., Sun, Y. & Sah, P. Neural circuits for a top-down control of fear and extinction. Psychopharmacology (Berl) 236, 313–320, doi:10.1007/s00213-018-5033-2 (2019).

23 Jovanovic, T., Kazama, A., Bachevalier, J. & Davis, M. Impaired safety signal learning may be a biomarker of PTSD. Neuropharmacology 62, 695–704, doi:10.1016/j.neuropharm.2011.02.023 (2012).

24 Acheson, D. T. et al. Conditioned fear and extinction learning performance and its association with psychiatric symptoms in active duty Marines. Psychoneuroendocrinology 51, 495–505, doi:10.1016/j.psyneuen.2014.09.030 (2015).

25 Jovanovic, T. et al. Impaired fear inhibition is a biomarker of PTSD but not depression. Depress Anxiety 27, 244–251, doi:10.1002/da.20663 (2010).

26 Grasser, L. R. Editorial: The Future of Safety Signal Learning as a Biomarker of Risk and Treatment Target for Trauma-Related Psychopathology in Youth. J Am Acad Child Adolesc Psychiatry 64, 772–774, doi:10.1016/j.jaac.2024.11.019 (2025).

27 Kausche, F. M., Carsten, H. P., Sobania, K. M. & Riesel, A. Fear and safety learning in anxiety- and stress-related disorders: An updated meta-analysis. Neurosci Biobehav Rev 169, 105983, doi:10.1016/j.neubiorev.2024.105983 (2025).

28 Foilb, A. R. & Christianson, J. P. in Neurobiology of Abnormal Emotion and Motivated Behaviors (eds Susan Sangha & Dan Foti) 204-222 (Academic Press, 2018).

29 Pollak, D. D. et al. An animal model of a behavioral intervention for depression. Neuron 60, 149–161, doi:10.1016/j.neuron.2008.07.041 (2008).

30 Kreutzmann, J. C., Jovanovic, T. & Fendt, M. Infralimbic cortex activity is required for the expression but not the acquisition of conditioned safety. Psychopharmacology (Berl) 237, 2161–2172, doi:10.1007/s00213-020-05527-7 (2020).

31 Laxmi, T. R., Stork, O. & Pape, H. C. Generalisation of conditioned fear and its behavioural expression in mice. Behav Brain Res 145, 89–98, doi:10.1016/s0166-4328(03)00101-3 (2003).

32 Hammond, L. J. A traditional demonstration of the active properties of Pavlovian inhibition using differential CER. Psychonomic Science 9, 65–66, doi:10.3758/BF03330761 (1967).

33 Rescorla, R. A. Conditioned inhibition of fear resulting from negative CS-US contingencies. J Comp Physiol Psychol 67, 504–509, doi:10.1037/h0027313 (1969).

34 Seligman, M. E. & Binik, Y. M. in Operant-pavlovian interactions 165–187 (Routledge, 1977).

35 Mombelli, E. et al. Auditory stimuli suppress contextual fear responses in safety learning independent of a possible safety meaning. Front Behav Neurosci 18, 1415047, doi:10.3389/fnbeh.2024.1415047 (2024).

36 LoLordo, V. M. & Fairless, J. L. Pavlovian conditioned inhibition: The literature since 1969. Information processing in animals: Conditioned inhibition, 1-49 (1985).

37 Rescorla, R. A. Pavlovian conditioned inhibition. Psychological bulletin 72, 77 (1969).

38 Takemoto, M. & Song, W. J. Cue-dependent safety and fear learning in a discriminative auditory fear conditioning paradigm in the mouse. Learn Mem 26, 284–290, doi:10.1101/lm.049577.119 (2019).

39 Christianson, J. P. et al. Safety signals mitigate the consequences of uncontrollable stress via a circuit involving the sensory insular cortex and bed nucleus of the stria terminalis. Biol Psychiatry 70, 458–464, doi:10.1016/j.biopsych.2011.04.004 (2011).

40 Baez, M. V., Cercato, M. C. & Jerusalinsky, D. A. NMDA Receptor Subunits Change after Synaptic Plasticity Induction and Learning and Memory Acquisition. Neural Plast 2018, 5093048, doi:10.1155/2018/5093048 (2018).

41 Johansen, J. P., Cain, C. K., Ostroff, L. E. & LeDoux, J. E. Molecular mechanisms of fear learning and memory. Cell 147, 509–524, doi:10.1016/j.cell.2011.10.009 (2011).

42 Ben Mamou, C., Gamache, K. & Nader, K. NMDA receptors are critical for unleashing consolidated auditory fear memories. Nat Neurosci 9, 1237–1239, doi:10.1038/nn1778 (2006).

43 Lee, H. & Kim, J. J. Amygdalar NMDA receptors are critical for new fear learning in previously fear-conditioned rats. J Neurosci 18, 8444–8454, doi:10.1523/JNEUROSCI.18-20-08444.1998 (1998).

44 Taft, C. E. & Turrigiano, G. G. PSD-95 promotes the stabilization of young synaptic contacts. Philos Trans R Soc Lond B Biol Sci 369, 20130134, doi:10.1098/rstb.2013.0134 (2014).

45 Ehrlich, I., Klein, M., Rumpel, S. & Malinow, R. PSD-95 is required for activity-driven synapse stabilization. Proc Natl Acad Sci U S A 104, 4176–4181, doi:10.1073/pnas.0609307104 (2007).

46 Bhattacharyya, S., Biou, V., Xu, W., Schlüter, O. & Malenka, R. C. A critical role for PSD-95/AKAP interactions in endocytosis of synaptic AMPA receptors. Nat Neurosci 12, 172–181, doi:10.1038/nn.2249 (2009).

47 de Oliveira Alvares, L. & Do-Monte, F. H. Understanding the dynamic and destiny of memories. Neurosci Biobehav Rev 125, 592–607, doi:10.1016/j.neubiorev.2021.03.009 (2021).

48 Laing, P. A. F., Vervliet, B., Dunsmoor, J. E. & Harrison, B. J. Pavlovian safety learning: An integrative theoretical review. Psychon Bull Rev 32, 176–202, doi:10.3758/s13423-024-02559-4 (2025).

49 Plas, S. L. et al. Neural circuits for the adaptive regulation of fear and extinction memory. Front Behav Neurosci 18, 1352797, doi:10.3389/fnbeh.2024.1352797 (2024).

50 LaBar, K. S. Neuroimaging of Fear Extinction. Curr Top Behav Neurosci 64, 79–101, doi:10.1007/7854_2023_429 (2023).

51 Krabbe, S., Grundemann, J. & Luthi, A. Amygdala Inhibitory Circuits Regulate Associative Fear Conditioning. Biol Psychiatry 83, 800–809, doi:10.1016/j.biopsych.2017.10.006 (2018).

52 Brockway, E. T., Simon, S. & Drew, M. R. Ventral hippocampal projections to infralimbic cortex and basolateral amygdala are differentially activated by contextual fear and extinction recall. Neurobiol Learn Mem 205, 107832, doi:10.1016/j.nlm.2023.107832 (2023).

53 Mavrych, V., Riyas, F. & Bolgova, O. The Role of Basolateral Amygdala and Medial Prefrontal Cortex in Fear: A Systematic Review. Cureus 17, e78198, doi:10.7759/cureus.78198 (2025).

54 Alexandra Kredlow, M., Fenster, R. J., Laurent, E. S., Ressler, K. J. & Phelps, E. A. Prefrontal cortex, amygdala, and threat processing: implications for PTSD. Neuropsychopharmacology 47, 247–259, doi:10.1038/s41386-021-01155-7 (2022).

55 Likhtik, E., Pelletier, J. G., Paz, R. & Pare, D. Prefrontal control of the amygdala. J Neurosci 25, 7429–7437, doi:10.1523/JNEUROSCI.2314-05.2005 (2005).

56 Herry, C. et al. Switching on and off fear by distinct neuronal circuits. Nature 454, 600–606, doi:10.1038/nature07166 (2008).

57 Zhang, X., Kim, J. & Tonegawa, S. Amygdala Reward Neurons Form and Store Fear Extinction Memory. Neuron 105, 1077–1093 e1077, doi:10.1016/j.neuron.2019.12.025 (2020).

58 Zhang, X., Flick, K., Rizzo, M., Pignatelli, M. & Tonegawa, S. Dopamine induces fear extinction by activating the reward-responding amygdala neurons. Proc Natl Acad Sci U S A 122, e2501331122, doi:10.1073/pnas.2501331122 (2025).

59 Sangha, S., Chadick, J. Z. & Janak, P. H. Safety encoding in the basal amygdala. J Neurosci 33, 3744–3751, doi:10.1523/JNEUROSCI.3302-12.2013 (2013).

60 Sangha, S. Plasticity of Fear and Safety Neurons of the Amygdala in Response to Fear Extinction. Front Behav Neurosci 9, 354, doi:10.3389/fnbeh.2015.00354 (2015).

61 Dunsmoor, J. E., Niv, Y., Daw, N. & Phelps, E. A. Rethinking Extinction. Neuron 88, 47–63, doi:10.1016/j.neuron.2015.09.028 (2015).

62 Marek, R., Strobel, C., Bredy, T. W. & Sah, P. The amygdala and medial prefrontal cortex: partners in the fear circuit. J Physiol 591, 2381–2391, doi:10.1113/jphysiol.2012.248575 (2013).

63 Spoormaker, V. I. et al. The neural correlates of negative prediction error signaling in human fear conditioning. Neuroimage 54, 2250–2256, doi:10.1016/j.neuroimage.2010.09.042 (2011).

64 Furlong, T. M., Cole, S., Hamlin, A. S. & McNally, G. P. The role of prefrontal cortex in predictive fear learning. Behav Neurosci 124, 574–586, doi:10.1037/a0020739 (2010).

65 Bouton, M. E., Maren, S. & McNally, G. P. Behavioral and Neurobiological Mechanisms of Pavlovian and Instrumental Extinction Learning. Physiol Rev 101, 611–681, doi:10.1152/physrev.00016.2020 (2021).

66 Junjiao, L. et al. Role of prediction error in destabilizing fear memories in retrieval extinction and its neural mechanisms. Cortex 121, 292–307, doi:10.1016/j.cortex.2019.09.003 (2019).

67 Dunsmoor, J. E. et al. Role of Human Ventromedial Prefrontal Cortex in Learning and Recall of Enhanced Extinction. J Neurosci 39, 3264–3276, doi:10.1523/JNEUROSCI.2713-18.2019 (2019).

68 Iordanova, M. D., Yau, J. O., McDannald, M. A. & Corbit, L. H. Neural substrates of appetitive and aversive prediction error. Neurosci Biobehav Rev 123, 337–351, doi:10.1016/j.neubiorev.2020.10.029 (2021).

69 Maren, S., Phan, K. L. & Liberzon, I. The contextual brain: implications for fear conditioning, extinction and psychopathology. Nat Rev Neurosci 14, 417–428, doi:10.1038/nrn3492 (2013).

70 Kalisch, R. et al. Context-dependent human extinction memory is mediated by a ventromedial prefrontal and hippocampal network. J Neurosci 26, 9503–9511, doi:10.1523/jneurosci.2021-06.2006 (2006).

71 Hughes, K. C. & Shin, L. M. Functional neuroimaging studies of post-traumatic stress disorder. Expert Rev Neurother 11, 275–285, doi:10.1586/ern.10.198 (2011).

72 Craske, M. G., Treanor, M., Conway, C. C., Zbozinek, T. & Vervliet, B. Maximizing exposure therapy: an inhibitory learning approach. Behav Res Ther 58, 10–23, doi:10.1016/j.brat.2014.04.006 (2014).

73 Benito, K. et al. Mechanisms of Change in Exposure Therapy for Anxiety and Related Disorders: A Research Agenda. Clin Psychol Sci 13, 687–719, doi:10.1177/21677026241240727 (2025).

74 Graner, J. L., Stjepanović, D. & LaBar, K. S. Extinction learning alters the neural representation of conditioned fear. Cogn Affect Behav Neurosci 20, 983–997, doi:10.3758/s13415-020-00814-4 (2020).

75 Goode, T. D., Kim, J. J. & Maren, S. Relapse of extinguished fear after exposure to a dangerous context is mitigated by testing in a safe context. Learn Mem 22, 170–178, doi:10.1101/lm.037028.114 (2015).

76 Meyer, H. C. & Lee, F. S. Intermixed safety cues facilitate extinction retention in adult and adolescent mice. Physiol Behav 271, 114336, doi:10.1016/j.physbeh.2023.114336 (2023).

77 Cooper, S. E. et al. Augmenting extinction with counterconditioning strengthens and sustains neural safety representations in PTSD. Translational Psychiatry 16, 303, doi:10.1038/s41398-026-03966-y (2026).

78 Kalinina, A., Krekhno, Z., Yee, J., Lehmann, H. & Fournier, N. M. Effect of repeated seizures on spatial exploration and immediate early gene expression in the hippocampus and dentate gyrus. IBRO Neurosci Rep 12, 73–80, doi:10.1016/j.ibneur.2021.12.008 (2022).

79 West, M. J., Slomianka, L. & Gundersen, H. J. Unbiased stereological estimation of the total number of neurons in the subdivisions of the rat hippocampus using the optical fractionator. Anat Rec 231, 482–497, doi:10.1002/ar.1092310411 (1991).

80 Paxinos, G. & Watson, C. *The Rat Brain in Stereotaxic Coordinates: Compact*. 7th edn, (Academic Press, 2017).

