## Supplementary Figures 1 and 2 for "Safety learning produces rapid fear suppression and distinct amygdala–prefrontal engagement"

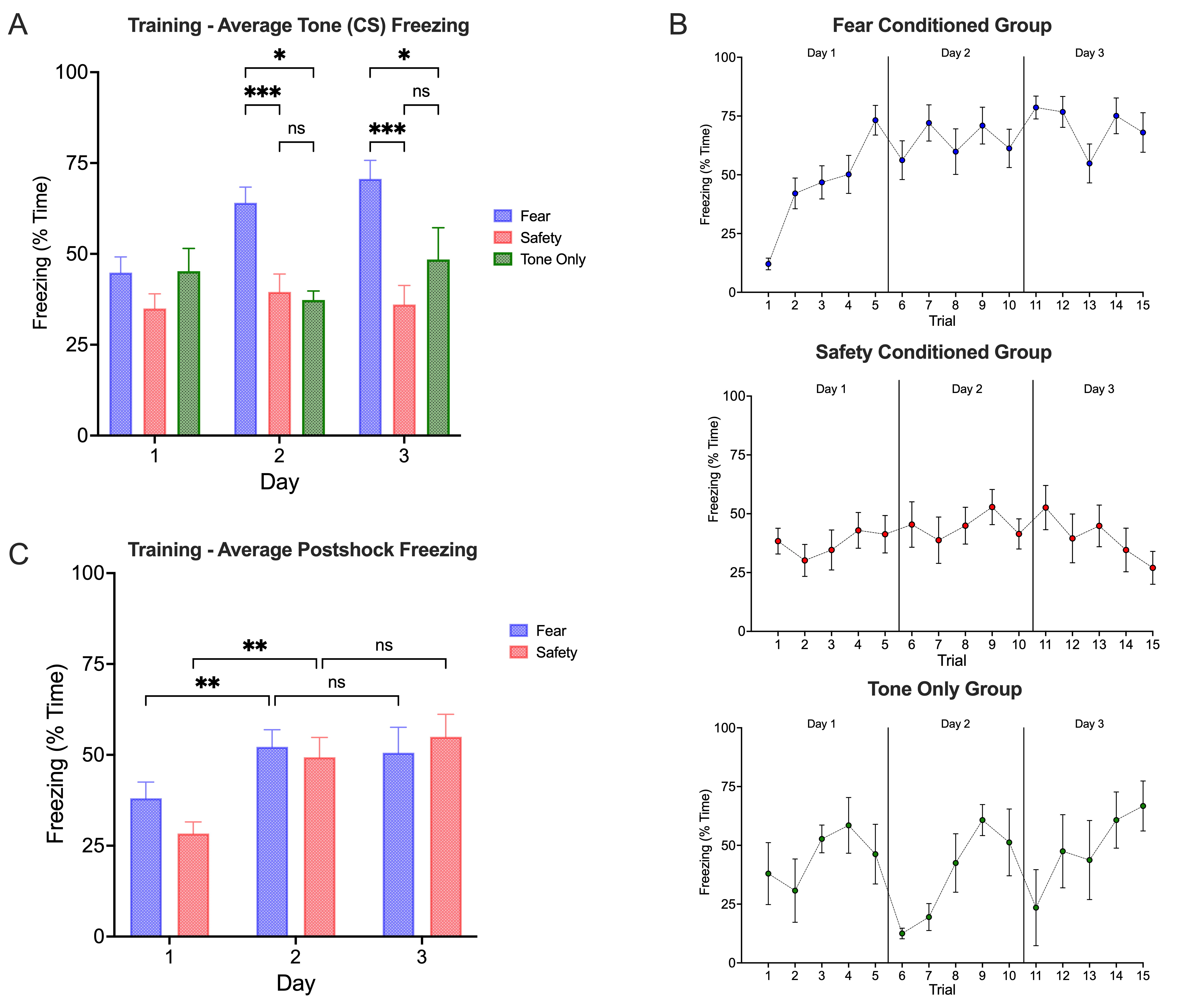

**Supplementary Fig. S1.** Tone-evoked and post-shock freezing during conditioning. (A) Mean percentage of time spent freezing during tone CS presentations across three days of conditioning. The numbers of CS and US presentations were matched between the fear and safety conditioned groups across three conditioning days, with one session conducted per day. Fear conditioned rats showed progressively greater CS-evoked freezing across conditioning, whereas safety conditioned rats showed lower CS-evoked freezing throughout conditioning. Tone-only controls showed intermediate levels of freezing that did not differ significantly from those of the safety conditioned group on Days 2 and 3. (B) Trial-by-trial freezing during CS presentations across all 15 tone trials for the three groups. (C) Mean post-shock freezing across conditioning days in the fear and safety conditioned groups. Post-shock freezing increased from Day 1 to Days 2 and 3, with no difference between the fear and safety conditioned groups. Data are presented as mean ± SEM. *p < .05, **p < .01, and ***p < .001; ns, not significant.

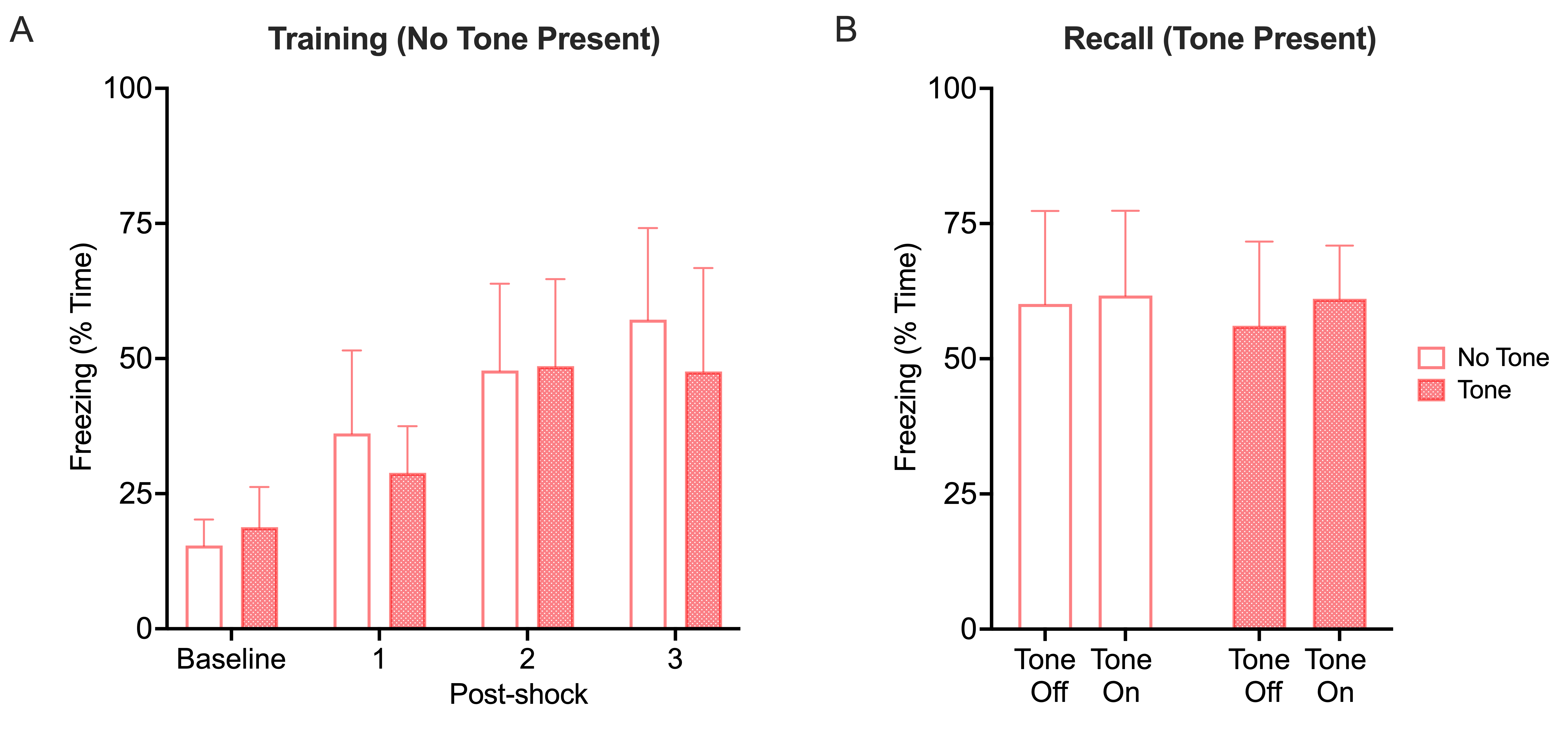
**Supplementary Fig. S2.** A novel tone presented during recall does not affect context freezing. (A) Mean percentage of time spent freezing during the baseline (pre-shock) and post-shock periods of contextual fear conditioning. (B) Mean percentage of time spent freezing during the recall test conducted 24 h later in rats tested either with or without presentation of a novel tone stimulus (20 s, 5 kHz, 80 dB). Presentation of the tone did not alter ongoing contextual freezing, with comparable freezing observed during the tone-off and tone-on periods. Freezing during these periods was also comparable to that observed across equivalent time periods in rats tested without the tone.
